# Activation of pro-survival autophagy by a small molecule promoting p62 oligomerisation

**DOI:** 10.1101/2025.05.27.656309

**Authors:** Johan Panek, Congsing Sun, Tetsushi Kataura, Edward Fielder, Laura Booth, Wyatt Yue, Gavin Richardson, Jóhannes Reynisson, Sovan Sarkar, Viktor I. Korolchuk

**Affiliations:** Biosciences Institute, Faculty of Medical Sciences, Newcastle University, Newcastle upon Tyne NE4 5PL, UK; Department of Cancer and Genomic Sciences, School of Medical Sciences, College of Medicine and Health, University of Birmingham, Birmingham B15 2TT, UK; Department of Neurology, Institute of Medicine, University of Tsukuba, Tsukuba, Ibaraki 305-8575, Japan; School of Allied Health Professions and Pharmacy, Keele University, Newcastle under Lyme, ST5 5BG, UK

**Keywords:** Autophagy, Niemann-Pick type C1 disease, Mitophagy, Oligomerisation, p62, ROS

## Abstract

Autophagy is a critical mechanism of cellular quality control, orchestrated by selective autophagy receptor (SAR) proteins. Pharmacologically enhancing the cargo-targeting capacity of SARs presents an attractive but underexplored strategy for the precise therapeutic activation of autophagy. Here, we characterise SQ-1, a small-molecule activator of autophagy that targets the prototypical SAR protein p62/SQSTM1 (sequestosome-1). We show that SQ-1 sensitises p62 to oxidation and promotes its disulphide-mediated oligomerisation in response to mitochondrial reactive oxygen species (ROS). This ROS-dependent activation of p62-mediated selective autophagy enhances the clearance of ROS-generating mitochondria and restores cell viability in models of Niemann-Pick type C1 (NPC1) disease, which is marked by impaired autophagic flux. In summary, the unique mode of action of SQ-1 enables self-regulated autophagy activation, offering a potential therapeutic strategy for lysosomal storage disorders and a broader spectrum of age-related diseases characterised by defective autophagy.

## Introduction

Macroautophagy (hereinafter autophagy) promotes proteostasis and longevity by degrading dysfunctional cellular components [1]. This homeostatic role of autophagy requires dedicated receptor proteins recognising damaged organelles and protein aggregates for the selective recruitment into autophagosomes, which deliver their cargo for degradation by lysosomes. p62 has been implicated in orchestrating several autophagy pathways by clustering ubiquitinated cargo and recruiting components of autophagy machinery for autophagosome assembly [2]. Oligomerisation of p62 is an essential step in promoting the avidity of its interactions with autophagy proteins such as the lipidated forms of Atg8/LC3 incorporated into the nascent autophagosome [3, 4]. The process of p62 oligomerisation is mediated by electrostatic intermolecular interactions between its Phox and Bem1 (PB1) domains and can be further facilitated by the oxidation of two cysteine residues (Cys105 and Cys113) forming disulphide-linked conjugates (DLC) [5–7]. Non-covalent PB1 domain-mediated interactions and DLC have been shown to promote the ability of p62 to induce autophagic degradation of ubiquitinated proteins and mitochondria (mitophagy) [5, 8].

Disruption of selective autophagy has been implicated in various diseases including neurodegeneration, cancer, and lysosomal storage disorders, underscoring the need for targeted strategies to restore autophagic function [9]. Several groups have recently developed small molecules targeting p62 and activating autophagy [8, 10, 11]. These small molecules are designed to mimic the N-terminal arginine protein degradation signal that is recognised by ZZ-type zinc finger domain of p62. By binding to the ZZ domain, they facilitate the formation of DLC and increase autophagy flux [6–8]. Our group has recently identified a novel p62-targeting small molecule STOCK1N which we re-named here to SQ-1 (a parent compound in “SeQuestra” series) [8]. SQ-1 robustly promotes the mitophagic clearance of dysfunctional mitochondria. This can mitigate and even reverse the development of cellular ageing phenotypes in primary human dermal fibroblasts [8].

At the same time, the molecular mechanisms by which p62-targeting small molecules promote DLC formation, selective autophagy, as well as their effect on cellular physiology remain poorly understood. It has been proposed that small molecule binding releases the interaction of the p62 ZZ domain with an unstructured linker region between PB1 and ZZ domains which contains Cys105 and Cys113 [11]. This can potentially expose these residues to the oxidising environment. Indeed, we demonstrated that oxidation-sensitive cysteine residues of p62 were essential for the ability of our p62-targeting small molecule to induce mitophagy [8]. However, the origin and the role of localised ROS triggering disulphide bond and DLC formation in this process has not been explored.

Here we show that residues Asp129 and Arg139 in p62 are required for the activation of autophagy by SQ-1, indicating these as the site of its binding within the ZZ domain of p62. Furthermore, both PB1 domain- and DLC-mediated oligomerisation of p62 are necessary for the autophagy-promoting effect of SQ-1 mediated by sensitisation of p62 to oxidation by mitochondrial ROS. By using cell models of Niemann-Pick type C1 disease associated with loss of function of the lysosomal cholesterol transporter NPC1 along with defective autophagic flux, we demonstrate that SQ-1 can reactivate dysfunctional autophagy/mitophagy, reduce mitochondrial load characterised by high levels of ROS, and rescue cell death of NPC1 patient-derived fibroblasts and neurons. Together, our study provides a mechanistic underpinning for the development of pharmacological interventions promoting selective autophagy in the context of lysosomal storage disease and potentially relevant to other neurodegenerative disorders.

## Results

### SQ-1 activates general and selective autophagy

Our previous study employed primary human dermal fibroblasts, commonly used as the model of cellular senescence, to demonstrate that activation of mitophagy by SQ-1 can potently rescue cellular ageing phenotypes [8]. To facilitate identification of molecular mechanisms underpinning the action of SQ-1, we employed penta-knockout (PentaKO) HeLa cell line with the loss of 5 SARs (p62, OPTN, NDP52, NBR1 and TAX1BP1) [12]. Stable expression of transgenic p62 constructs in this cell line allowed us to investigate p62-specific effects of SQ-1 in the absence of compounding factors mediated by the related SAR family members.

To validate previous findings in this model, PentaKO cells expressing Halo-tagged wild-type p62 construct were labelled with Halo ligand (Janelia Fluor 646) and treated for 5 hours with 40 µM SQ-1 or 50 nM rapamycin, a potent autophagy inducer used as a control [13, 14]. Both compound treatments strongly increased the formation of p62 puncta, which, in the presence of the mRFP-GFP-LC3 autophagy reporter, colocalised with both GFP^+^ and RFP^+^ structures (Fig. 1A, B) [15]. This indicated that, similar to rapamycin, SQ-1 induces assembly of p62 into autophagic vesicles. However, in contrast to rapamycin, the effect of SQ-1 was not mediated by the suppression of mechanistic Target Of Rapamycin Complex 1 (mTORC1) activity (Suppl. Fig. 1A) [13].

**Figure 1.**
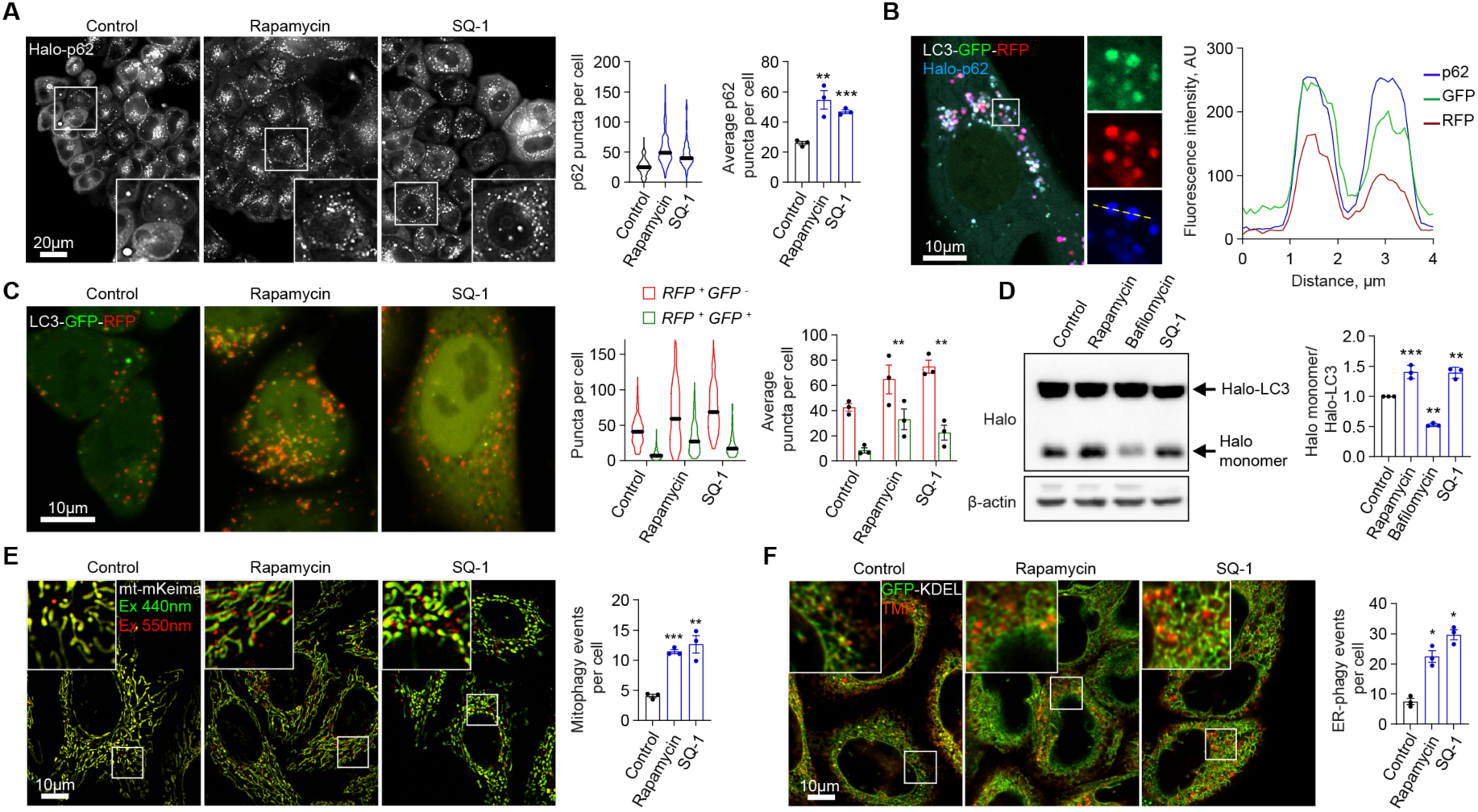
Small molecule SQ-1 promotes inclusion of p62 into autophagosomes and activates selective autophagy. **A,** Fluorescence microscopy images and quantification of p62 puncta in PentaKO HeLa cells expressing Halo-p62 labelled with Halo ligand (Janelia Fluor 646) and treated with 50 nM rapamycin or 40 µM SQ-1 for 5 h. **B**, Confocal microscopy images of PentaKO HeLa cells expressing mRFP-GFP-LC3 autophagy reporter and Halo-p62 labelled with Halo ligand and treated with 40 µM SQ-1 for 5 h. The plot is showing the intensity values for p62, GFP and RFP signal over the transect indicated by the yellow line. **C**, Fluorescence microscopy images and quantification of autophagosomes and autolysosomes in HeLa cells expressing mRFP-GFP-LC3 cultured in DMEM supplemented with 50 nM rapamycin or 10 µM SQ-1 for 24 h. **D**, Immunoblot analyses and quantification of Halo monomer accumulation in HeLa cells expressing Halo-LC3 treated with 50 nM rapamycin, 400 nM bafilomycin A1, or 40 µM SQ-1 in the presence of HaloTag ligand for 5 h. **E**, Fluorescence microscopy images and quantification of mitophagy in HeLa cells expressing mt-mKeima reporter cultured in galactose medium supplemented with 50 nM rapamycin or 10 µM SQ-1 for 24 h. **F**, Fluorescence microscopy images and quantification of ER-phagy in HeLa cells expressing Halo-GFP-KDEL reporter cultured in DMEM medium supplemented with 50 nM rapamycin or 10 µM SQ-1 and 1 µM Halo TMR ligand for 5 h. Data are mean ± SEM of n = 3 biological replicates. P values were calculated by one-way ANOVA followed by multiple comparisons with Dunnett’s test (A, C, E, F) or two-way ANOVA followed by multiple comparisons with Dunnett’s test (D). *P< 0.05; **P<0.01; ***P<0.001; ns (non-significant).

To compare the effect of SQ-1 and rapamycin on the activation of autophagic flux, mRFP-GFP-LC3 expressing HeLa cells were treated with the compounds for 24 hours and analysed by fluorescence microscopy to quantify the number of autophagosomes (RFP^+^, GFP^+^) and autolysosomes (RFP^+^, GFP^-^) [13]. Both treatments significantly increased the levels of both early- and late-stage autophagic vesicles which demonstrates a strong induction of the entire autophagy pathway (Fig. 1C). An orthogonal immunoblotting assay using cells expressing Halo-LC3 reporter confirmed similar induction of autophagy flux in response to SQ-1 and rapamycin treatment, whilst inhibition of autophagy by bafilomycin A1 validated the specificity of the assay (Fig. 1D) [14]. To investigate the effect of SQ-1 and rapamycin on selective forms of autophagy, we employed HeLa cells expressing a mitophagy reporter mt-mKeima and an endoplasmic reticulum autophagy (ER-phagy) reporter Halo-GFP-KDEL [12, 14]. Both types of selective autophagy were strongly induced by the treatments (Fig. 1E, F) indicating that SQ-1 is a potent activator of general and selective autophagy pathways.

### SQ-1 activates autophagy in a p62-dependent manner

Next, we used PentaKO HeLa cells expressing mRFP-GFP-LC3 in the absence or presence of Halo-p62 constructs to investigate the requirement for p62 during autophagy activation by SQ-1 and rapamycin. SQ-1 increased the formation of autophagosomes or autolysosomes in cells expressing wild-type Halo-tagged p62, but not empty Halo vector, indicating that p62 is both necessary and sufficient for the induction of autophagy flux in response to the small molecule (Fig. 2A). At the same time, rapamycin significantly increased autophagy in a p62-independent manner consistent with its established mechanism of autophagy activation [15]. Cells expressing p62 mutants defective for DLC formation (C105A,C113A-p62) or PB1 domain oligomerisation (K7A,D69A-p62) phenocopied cells carrying empty Halo construct. In these cells, rapamycin, but not SQ-1, could activate autophagy (Fig. 2A) [5, 8]. Together, these data demonstrate that induction of autophagy by SQ-1 is strictly dependent on p62 and requires its ability to oligomerise, both via disulphide bonds and PB1 domain interactions.

**Figure 2.**
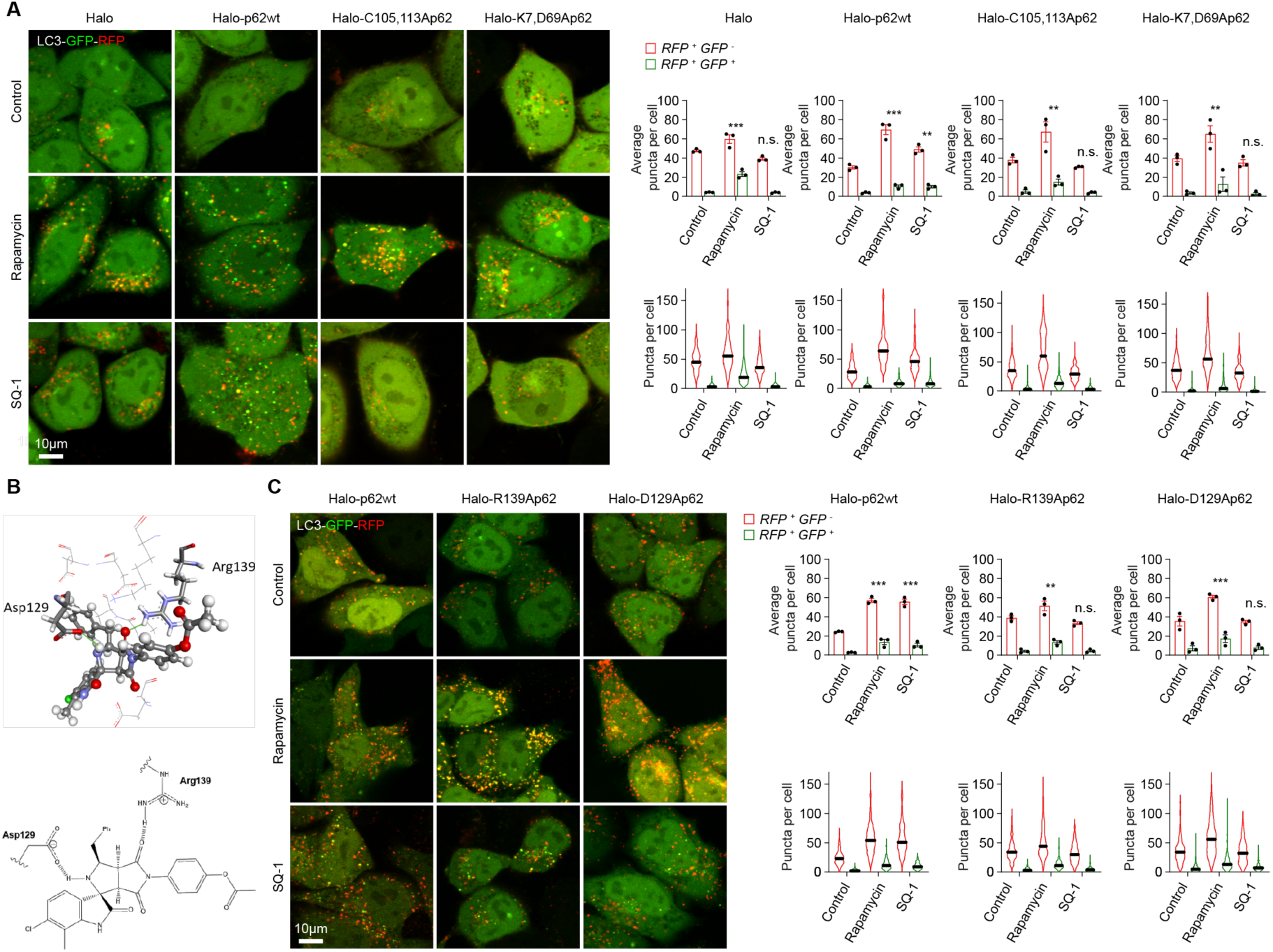
Induction of autophagy by SQ-1 is dependent upon ZZ domain and oligomerisation of p62. **A**, Fluorescence microscopy images and quantification of autophagosomes and autolysosomes in PentaKO HeLa cells expressing mRFP-GFP-LC3 and Halo-p62wt, Halo-C105A,C113Ap62 or Halo-K7A,D69Ap62 cultured in DMEM supplemented with 50 nM rapamycin or 10 µM SQ-1 for 24 h. **B**, The docked pose of SQ-1 (ball-and-stick format) in the binding site of p62 as predicted by the ASP scoring function. H-boding is predicted between the carboxylic side chain of Asp129 and the pyrrolidine amine group of the central ring scaffold of the ligand (2.1 Å) as well as Arg139’s guanidine side chain with one of the carbonyl groups of the succinimide ring (1.7 Å). The adjacent amino acids (< 5 Å), buttressing the ligand, are shown as lines. The amino acids’ hydrogens are not shown for clarity. Below is the two-dimensional structure of SQ-1 with the H-bonding interactions shown. **C**, Fluorescence microscopy images and quantification of autophagosomes and autolysosomes in PentaKO HeLa cells expressing mRFP-GFP-LC3 and Halo-p62wt, Halo-R139Ap62 or Halo-D129Ap62 cultured in DMEM supplemented with 50 nM rapamycin or 10 µM SQ-1 for 24 h. Data are mean ± SEM of n = 3 biological replicates. P values were calculated by two-way ANOVA followed by multiple comparisons with Dunnett’s test (A, C). **P<0.01; ***P<0.001; ns (non-significant).

Our modelling of SQ-1 complex with ZZ domain of p62 indicated that, similar to other synthetic p62 ligands, amino acid residues Asp129 and Arg139 become engaged in critical electrostatic interactions with the small molecule (Fig. 2B) [7, 11]. p62 constructs carrying Ala substitutions of these residues were generated and expressed together with the mRFP-GFP-LC3 reporter in PentaKO HeLa. As shown in Fig. 2C, Asp129 and Arg139 were found to be essential for the ability of SQ-1 (but not rapamycin) to induce autophagy flux. We conclude that binding of SQ-1 to the specific pocket within ZZ domain containing Asp129 and Arg139, as well as oligomerisation of p62, are essential elements of the mechanism underpinning the activation of autophagy by SQ-1.

### Activation of autophagy by SQ-1 requires ROS

SQ-1, as well as p62-targeting small molecules developed by others, promote the formation of p62 DLC that are essential for the ability of SQ-1 to induce autophagy [7, 8]. However, the mechanism by which the small molecules promote oxidation of redox-reactive cysteine residues triggering disulphide formation remains unknown. Our density functional theory (DFT) calculations indicated that SQ-1 is redox stable. The ionisation potential (one-electron oxidation) of SQ-1 is 7.52 eV with 95% of drugs lying in the 6.0 – 9.0 eV range; the electron affinity (one-electron reduction) is -1.16 eV with drugs ranging from -1.5 to 2.0 eV (95% confidence interval) [16]. SQ-1 is in the middle of the ionisation range but relatively low for the electron affinity albeit still within the reported range of known drugs (Suppl. Table 1). The low electron affinity can most easily be explained by the ester group occupying the *para* position on one of the phenyl rings (Fig. 2B). Therefore, oxidation of p62 is unlikely to be caused by SQ-1 directly.

We investigated if SQ-1 may instead sensitise p62 to oxidation by intracellular ROS. We exposed HeLa cells to a short bout of low concentration hydrogen peroxide (100 µM, 10 min) following 18 h pre-treatment of cells with SQ-1 or vehicle. Immunoblot analyses in non-reducing conditions, and immunofluorescence detection of intracellular puncta were used to ascertain the formation of DLC and self-assembly of p62, respectively [5, 8]. SQ-1 pre-treatment in the absence of exogenous H_2_O_2_ produced small but not statistically significant increase in DLC and p62 puncta formation (Fig. 3A, B). This is in contrast to the data presented in Fig. 1A where higher concentration was used in a shorter treatment. Importantly, in cells pre-treated with SQ-1 the formation of p62 DLC and microscopically observable puncta was significantly higher when cells were given a short boost of hydrogen peroxide (Fig. 3A, B). These data are consistent with the notion of p62 sensitisation to oxidation by SQ-1 binding.

**Figure 3.**
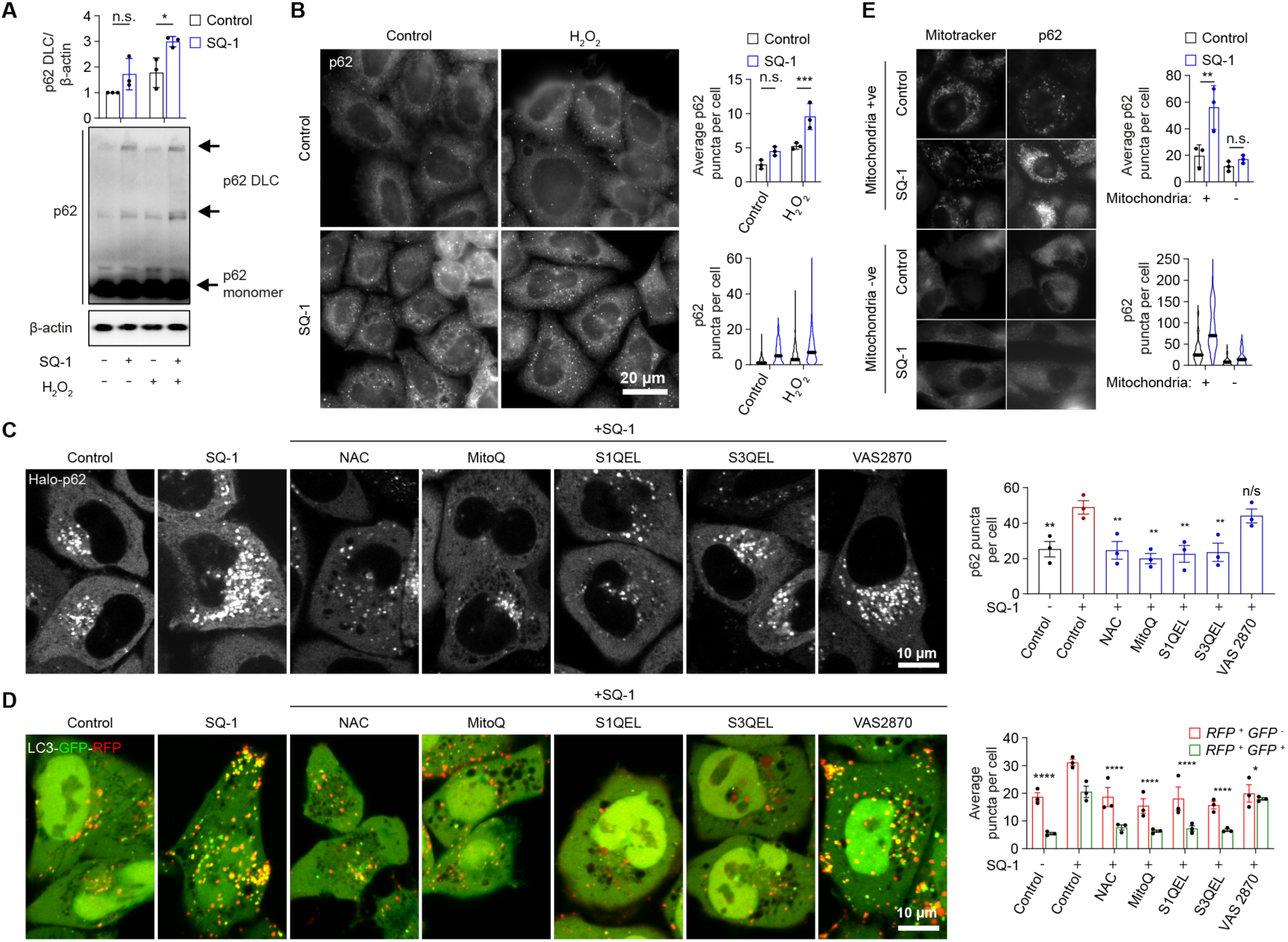
SQ-1 induces autophagy by sensitising p62 to oxidation by mitochondria-derived ROS. **A**, Immunoblot analyses in non-reducing condition and quantification of p62 DLCs accumulation in HeLa cells treated with 30 µM SQ-1 for 18 h followed by H_2_O_2_ at 100 µM for 10 min. **B,** Immunofluorescence microscopy images and quantification of p62 puncta accumulation in HeLa cells treated as in A. **C**, Fluorescence microscopy images and quantification of p62 puncta in PentaKO HeLa cells expressing Halo-p62 labelled with Halo ligand (Janelia Fluor 646) and treated with 40 µM SQ-1 and 1 mM NAC, 500 nM MitoQ, 10 µM S1QEL, 10 µM S3QEL or 10 µM VAS 2870 for 5 h. **D**, Confocal fluorescence microscopy images and quantification of autophagosomes and autolysosomes in HeLa cells expressing mRFP-GFP-LC3 treated as in C. **E**, Fluorescence microscopy images and quantification of p62 puncta in mitochondria-retaining (mitochondria +ve) and mitochondria-deficient (mitochondria -ve) cells labelled with Halo ligand (Janelia Fluor 646) ±40 µM SQ-1 in PentaKO HeLa cells expressing Halo-p62 (previously transiently transfected to over-express YFP-Parkin and then treated with 2×24 hours of 10 µM FCCP). Data are mean ± SEM of n = 3 biological replicates. P values were calculated by one-way ANOVA followed by multiple comparisons with Dunnett’s test (C) or two-way ANOVA followed by multiple comparisons with Dunnett’s (D) and Šidák’s test (A, B, E). *P< 0.05; **P<0.01; ***P<0.001; ns (non-significant).

Next, we investigated the requirement for, and the source of, the endogenously generated ROS triggering p62 oligomerisation and autophagy induction in response to SQ-1 treatment. We tested several established chemical compounds: N-Acetyl-L-cysteine (NAC) as a general antioxidant, mitochondria-targeted superoxide scavenger Mito Q10 (MitoQ), inhibitors of superoxide generation by mitochondrial Complex I (S1QEL) or Complex III (S3QEL), and an inhibitor of cytoplasmic NADPH oxidase VAS2870 [17, 18]. Increased formation of p62 puncta, autophagosomes and autolysosomes induced by SQ-1 treatment was completely suppressed by NAC or mitochondria-targeted molecules. In contrast, inhibitor of NADPH oxidase had little effect on SQ-1-induced p62 puncta formation and autophagosome/autolysosome numbers (Fig. 3C, D). The effect of antioxidants was specific to SQ-1 response as formation of p62 puncta and autophagosome/autolysosome formation was not significantly affected in basal conditions (Suppl. Fig. 1B, C).

Given that mitochondrial-targeted, but not cytoplasmic NAPDH oxidase-targeted, antioxidants negated the SQ-1-induced increase in p62 puncta, we wanted to test whether mitochondria themselves were required for the effect of SQ-1. We transiently transfected p62 expressing PentaKO HeLa cells with YFP-Parkin and then induced mitophagy with high doses of carbonyl cyanide-p-trifluoromethoxyphenylhydrazone (FCCP), an established method to generate cells without mitochondria [19, 20]. By doing this, we were able to deplete mitochondria from the transfected sub-population of cells (Fig. 3E). p62 puncta were predominantly present in cells retaining morphologically distinct mitochondria compared to mitochondria-depleted cells whilst SQ-1 treatment amplified p62 puncta formation specifically in cells retaining mitochondria (Fig. 3E). We conclude that physiological levels of ROS, particularly those generated during respiration by the mitochondrial electron transport chain, are essential for SQ-1-induced p62 oligomerisation and autophagy activation.

### SQ-1 rescues cellular deficits due to NPC1 deficiency

Autophagy flux is disrupted in cellular models of NPC1 disease [13, 21, 22]. In actively respiring cells with the loss/mutation of NPC1 protein, such as mouse embryonic fibroblasts (MEFs) cultured in galactose medium, or upon differentiation of patient-derived induced pluripotent stem cells (iPSC) into neurons, the loss of mitochondrial quality control by autophagy results in an increased production of ROS and cell death [13, 17]. We have shown that autophagy inducers, including rapamycin, celecoxib and memantine, can restore autophagy/mitophagy flux in NPC1 deficient cells, reduce accumulation of mitochondria with high ROS levels, and rescue cell survival [13, 21]. Similar to rapamycin, SQ-1 treatment resulted in the rescue of these phenotypes in *Npc1^-/-^* MEFs cultured in galactose medium (Fig. 4A-B, Suppl. Fig. 1D) [13]. However, unlike rapamycin which rescues cell death in the absence of autophagy [13], SQ-1 did not rescue survival of autophagy-deficient *Atg5^-/-^*MEFs cultured in galactose medium. These data indicate that the effect of SQ-1 on cell survival is strictly dependent on autophagy (Fig. 4C).

**Figure 4.**
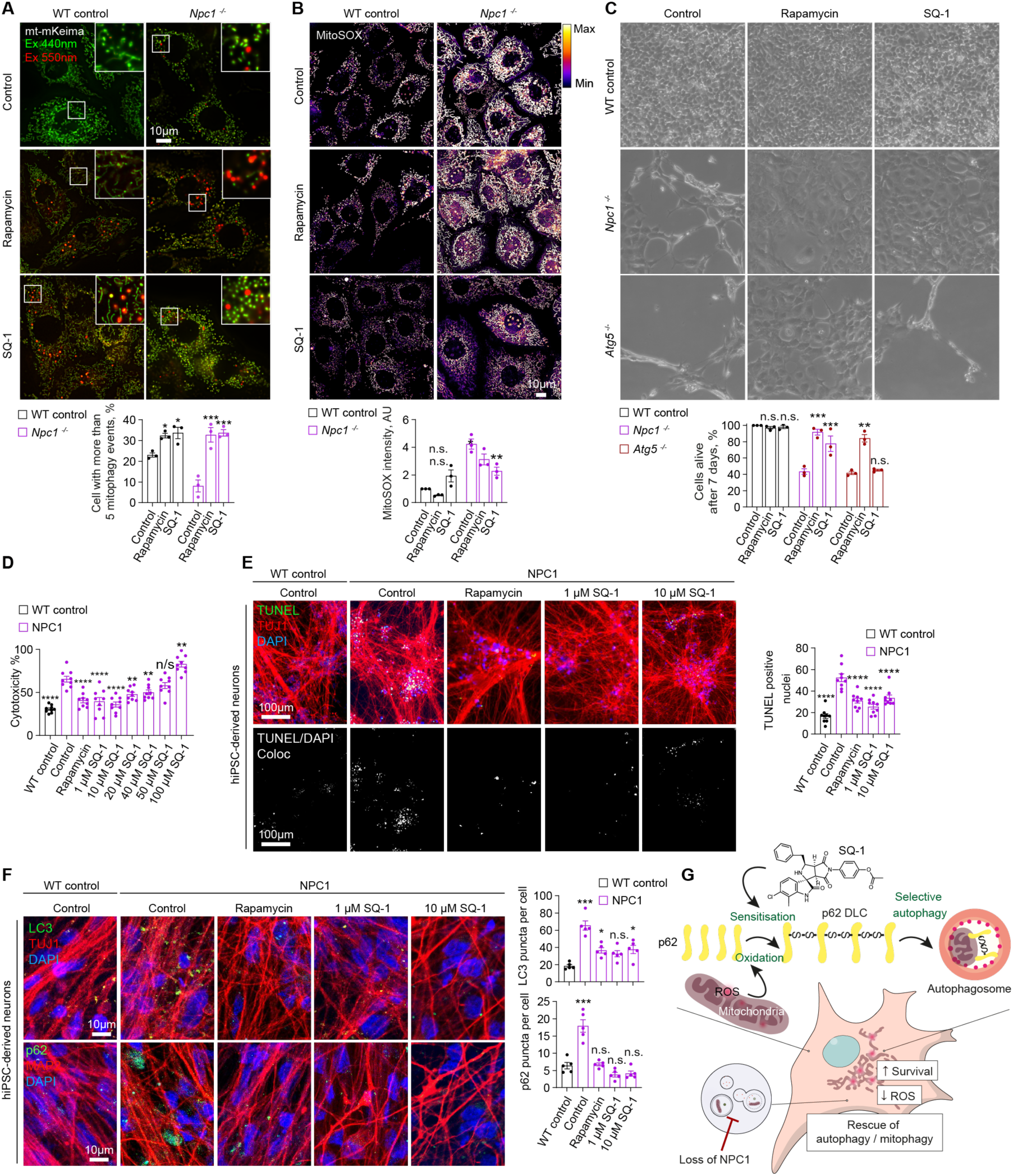
SQ-1 rescues survival of NPC1 cells by activating autophagy/mitophagy. **A,** Fluorescence microscopy images and quantification of mitophagy in *Npc1*^+/+^ and *Npc1*^-/-^ MEFs expressing mt-mKeima cultured for 24 h in galactose medium supplemented with 50 nM rapamycin or 10 µM SQ-1. **B**, Fluorescence microscopy images and quantification of MitoSOX staining of *Npc1*^+/+^ and *Npc1*^-/-^ MEFs cultured for 24 h in galactose medium supplemented with 50 nM rapamycin or 10 µM SQ-1. **C**, Phase-contrast images and cytotoxicity assay in *Npc1^+/+^* and *Npc1^-/-^* MEFs cultured in galactose medium supplemented with 50 nM rapamycin or 10 µM SQ-1 for 72 h (phase contrast image) or 144 h (cytotoxicity assay). **D**, Cytotoxicity assay in control and NPC1 patient iPSC-derived cortical neurons after 4 weeks of neuronal differentiation, where neurons were treated with 50 nM rapamycin or 1-100 µM SQ-1 for 6 days. **E**, Fluorescence images and quantification of TUNEL^+^ apoptotic nuclei in TUJ1^+^ cells treated as in D. **F**, Fluorescence images and quantification of LC3B and p62 puncta in TUJ1^+^ treated as in D. **G**, Schematic representation of the proposed mechanism by which SQ-1 sensitises p62 to mitochondria-generated ROS, leading to p62 oligomer formation, selective autophagy/mitophagy rescue, reduction in mitochondria with elevated ROS levels, and rescue of cell death. Data are mean ± SEM of n = 3 biological replicates. P values were calculated by one-way ANOVA followed by multiple comparisons with Dunnett’s test (A) or two-way ANOVA followed by multiple comparisons with Dunnett’s test (B, C, D). *P< 0.05; **P<0.01; ***P<0.001; ns (non-significant).

Next, we utilised NPC1 patient iPSC-derived neuronal cells, which are disease-relevant cellular platforms widely used to evaluate the efficacy of candidate drugs. NPC1 neurons exhibit increased cell death alongside a block in autophagic flux, involving accumulation of autophagosomes and p62, compared to the control neurons [13, 17, 23]. We tested a range of SQ-1 concentrations and found that doses between 1 and 20 µM resulted in the rescue of cell death similar to rapamycin. At higher concentrations the cytoprotective effect was abolished, likely due to compound cytotoxicity within our extended treatment protocol (Fig. 4D). We therefore selected the lowest two concentrations (1 and 10 µM) to further investigate their effect on autophagy and cell death in neurons. Assessment of TUNEL staining for apoptotic nuclei in TUJ1-positive neuronal cells indicated that SQ-1, similar to rapamycin, effectively rescued cell death of NPC1 neurons (Fig. 4E) [13, 17]. This cytoprotective effect was associated with reduced accumulation of autophagosomal markers LC3 and p62 in NPC1 neurons, demonstrating the restoration of autophagic flux (Fig. 4F).

Based on these data, we propose that by sensitising p62 to oxidation by mitochondria-derived ROS, SQ-1 enhances the flux through p62-dependent selective autophagy pathways. This reduces the build-up of autophagosomes and accumulation of damaged cellular components, including mitochondria, characterised by elevated ROS levels in cells with NPC1 deficiency. Ultimately, the restoration of cellular quality control mechanisms results in the rescue of cell death caused by NPC1 deficiency (Fig. 4G).

## Discussion

Selective autophagy acts as an important quality control mechanism by eliminating dysfunctional cellular components [1]. As such, pharmacologically enhancing the activity of selective autophagy pathways may potentially offer novel therapeutic opportunities in a broad range of human diseases associated with an accumulation of damaged organelles such as mitochondria generating high levels of ROS. Targeting SARs, the molecular machines that recognise the cargo and directly interact with the core autophagy machinery, presents a unique approach to upregulating selective autophagy. By targeting p62 we can induce autophagy in a specific and precise manner. This is in contrast to the majority of established autophagy activators, which modulate upstream signalling cascades and inevitably result in pleiotropic effects on cell biology [13, 24].

Our current study provides a mechanistic underpinning for the high specificity of p62 targeting ligands, such as SQ-1. The small molecule potentiates the response of p62 to the physiologically occurring changes in redox homeostasis, likely triggered by localised elevated levels of ROS produced within the mitochondrial network. It can be hypothesised that by sensitising p62 to the ROS signal and promoting its disulphide formation and oligomerisation, SQ-1 facilitates targeting of the potentially damaged mitochondrial component for p62-dependent mitophagy. One open question is the exact source and the nature of ROS promoting p62 DLC formation. The most likely explanation is that superoxide produced by mitochondrial complex I and III is dismutated into H_2_O_2_ which permeates into cytoplasm where it oxidises p62 [18]. However, the communication of mitochondria with the autophagy machinery is likely to be complex and dependent on different mitochondrial states. For example, mitochondrial depolarisation and increased permeability associated with damage may allow for the leakage of superoxide into cytoplasm [18]. Instead, high levels of carbon dioxide and ROS produced by metabolically active mitochondria will result in the formation of peroxymonocarbonate, a potent oxidising agent which may similarly act as a localised cue for p62 DLC formation, albeit in response to the different mitochondrial state [25].

It will be particularly important to investigate oligomerisation of p62 and the activation of autophagosome formation at the sub-mitochondrial level, in order to understand specific characteristics of mitochondria being targeted by p62 dependent mitophagy in response to SQ-1 treatment. The same arguments apply to the mechanisms by which p62 oligomerisation/DLC formation may play the role in selective autophagy pathways other than mitophagy. For example, our data indicate that SQ-1 treatment promotes ER-phagy, however it remains to be investigated if this is a bystander effect of the p62-dependent autophagy activation, or whether as suggested by others p62 is directly involved in the quality control of ER, alone or as the component of the mitochondria-ER contact sites [26].

Another important question pertaining to the mechanism of the p62-targeting small molecules is how binding the ZZ domain increases the protein’s propensity to disulphide formation. Recently a p62 ligand dusquetide has been proposed to bind two ZZ domains in the trans manner, effectively acting as a molecular staple bringing together two p62 monomers [10]. However, our modelling studies did not identify potential binding of SQ-1 with the second p62 ZZ domain moiety. Instead, the model where binding of the ligand releases the linker region containing redox-regulated cysteine residues from its interaction with the ZZ domain thus allowing for the formation of intermolecular disulphide bonds appears to be the most likely hypothesis [11]. Overall, our site-directed mutagenesis of the p62 ZZ domain supports the *in silico* model of SQ-1 binding to its docking site, as well as the requirement for the disulphide formation during ligand-induced autophagy. However, this model would require experimental testing, including the rigorous investigation of the direct binding of SQ-1 to p62 using structural biology and biophysics approaches as part of the effort to develop p62-targeting ligands into drug-like entities.

Importantly, our current data provide strong evidence for the cytoprotective effect of SQ-1 in cell models of NPC1, including patient iPSC-derived neurons. These data set the ground for future medicinal chemistry work and *in vivo* testing of p62-specific ligands. What is clear however, that should these efforts be successful, pharmacological interventions meaningfully improving cellular quality control mechanisms could be of significant and broad relevance to medicine, from rare forms of lysosomal storage diseases to slowing down the ageing process itself.

## Materials and Methods

### Culture of mammalian cell models

Immortalized *Npc1*^+/+^ and *Npc1^-/-^* MEFs (gift from Peter Lobel); *Atg5^+/+^* and *Atg5^-/-^* MEFs (gift from Noboru Mizushima) [27]; wild type and PentaKO HeLa cells [12] were maintained in DMEM supplemented with 10% foetal bovine serum (FBS), 100 U/mL penicillin/streptomycin and 2 mM L-glutamine (all from Sigma-Aldrich) at 37 °C, and 5% CO_2_ in a humidified incubator. 293FT cells (R70007, Invitrogen) were cultured as above in medium supplemented with 1X MEM non-essential amino acids (Gibco).

### iPSC culture and neuronal differentiation

Human induced pluripotent stem cell (iPSC) lines, control and NPC1 iPSC lines were cultured as previously described [23, 28]. The human iPSC lines were cultured on inactivated mouse embryonic fibroblast (MEF) feeder layer in hESC medium comprised of DMEM/F-12, 5% KnockOut Serum Replacement, 1% L-glutamine, 1% non-essential amino acids, 1% penicillin/streptomycin, 4 ng/mL human recombinant basic fibroblast growth factor (bFGF) (all from Gibco), 15% FBS (HyClone) and 0.1 mM β-mercaptoethanol (Sigma-Aldrich) and maintained in a humidified incubator with 5% CO_2_ and 5% O_2_ at 37 °C. For experimentation, the iPSCs were cultured feeder-free on Geltrex basement membrane matrix in StemFlex Basal Medium supplemented with StemFlex 10X Supplement (all from Gibco).

Neural stem cells (NSCs) were differentiated from iPSCs, as described previously [29, 30]. NSCs were cultured on Poly-L-ornithine and Laminin (PO-L) (Sigma-Aldrich) coated plates or flasks in N2B27 medium comprised of DMEM/F-12 and Neurobasal medium in 1:1 ratio, 1% N-2 supplement, 2% B-27 supplement, 1% penicillin/streptomycin (all from Gibco), 0.1% β-mercaptoethanol (Sigma-Aldrich) supplemented with 10 ng/mL FGF-2 (Miltenyi Biotec) and 10 ng/mL EGF (PeproTech), and were maintained in a humidified incubator with 5% CO_2_ at 37 °C. NSCs were passaged twice a week with 0.05% Trypsin-EDTA (Gibco), and the medium was changed on alternate days.

Neuronal differentiation of human iPSC-derived NSCs was carried out as described previously [29, 30]. The NSCs were seeded as above on PO-L coated plates in N2B27 medium without FGF-2 and EGF. At day 4 of neuronal differentiation, cells were treated with 10 µM DAPT (Tocris) to prevent cell proliferation. The N2B27 medium (without FGF-2 and EGF) was changed every 2 days and neuronal differentiation was carried out for 4 weeks. The neurons generated *in vitro* were cortical in nature [29, 30].

The control and NPC1 patient-derived iPSCs were originally generated in the lab of Rudolf Jaenisch at the Whitehead Institute for Biomedical Research. These cell lines were used for this study in the lab of Sovan Sarkar at the University of Birmingham under material transfer agreements, UBMTA 15-0593 and UBMTA 15-0594. All experiments were performed in accordance with ISSCR and institutional guidelines and regulations.

### Generation of stable cell lines

Generation of cells stably expressing YFP-Parkin, mt-mKeima, Halo-LC3, Halo-GFP-KDEL, mRFP-GFP-LC3, or the Halo-p62 wild-type and mutant (K7A,D69A-p62, C105A,C113A-p62, D129A-p62, R139A-p62) constructs were achieved by packaging retroviruses or lentiviruses in the HEK293FT (293FT) cell line. Cells were seeded in a 10 cm dish (6.0×10^6^ cells/10 mL/dish) in antibiotic-free culture medium. The next day, cells were transfected with plasmids containing the packaging gag/pol (retrovirus) or psPAX2 (lentivirus) and envelope pCMV-VSV-G genes, and the YFP-Parkin-IRES-zeo (gift from Douglas Green and Stephen Tait, Addgene, 61728) [31], pCHAC-mt-mKeima (gift from Richard Youle, Addgene, 72342) [12], pMRX-IP-HaloTag7-LC3 and pMRX-IB-HaloTag7-mGFP-KDEL (gift from Noboru Mizushima, Addgene, 184901) [32] or Halo-p62 constructs (ordered from Vector Builders) using Lipofectamine 3000. Following overnight transfection, the medium was replaced with fresh antibiotic-free medium. After 48 h virus containing medium was collected and concentrated using LentiX concentrator reagent (Takara Bio). The different cell lines were then transduced using concentrated virus and selected after 24 h using the appropriate selection antibiotic: Cells expressing mRFP-GFP-LC3, Halo-LC3 and the different Halo-p62 mutants were selected using 8 μg/ml of puromycin (Invitrogen). Cells expressing Halo-GFP-KDEL were selected using 4 μg/ml of blasticidin. Following selection, PentaKO HeLa cells transduced with Halo-p62 mutant constructs were labelled with Janelia Fluor 646 Halo ligand and sorted by Fluorescence-Activated Cell Sorting (FACS, BD FACSAria Fusion) to select cells with p62 expression level matching *wt* Halo-p62. To generate cell lines expressing both Halo-p62 mutant construct and mRFP-GFP-LC3 reporter, previously sorted cells were transduced second time with mRFP-GFP-LC3 and GFP expressing cells were selected by FACS (BD FACSAria Fusion).

### p62 puncta formation assay

PentaKO HeLa cells transduced with either wild type or mutant Halo tagged p62 were seeded into 24-well glass bottom plates (Greiner). 24 hours later, cells were labelled with 40 nM Janelia Fluor 646 Halo ligand (GA1120, Promega) and then incubated for 5 h with following compounds: 50 nM rapamycin (R8781-200UL, Sigma-Aldrich), 400 nM bafilomycin A1 (BML-CM110-0100, Enzo Life Sciences), 40 µM SQ-1 (STOCK1N-57534, InterBioScreen Ltd), 1 mM N-Acetyl-L-cysteine (A7250, Sigma-Aldrich), 500 nM MitoQ10 (S8978-SEL, Stratech Scientific Ltd), 10 µM S1QEL (F2068-0013, Life Chemicals), 10 µM S3QEL (SML1554, Sigma-Aldrich) and 10 µM VAS 2870 (BML-EI395-0010, Enzo Life Sciences). Live cells were then imaged on an inverted DMi8 microscope (Leica) with a Plan-Apochromat 63x/1.40 oil immersion lens. The number of puncta per cell was quantified by outlining single cells as regions of interest using Cellpose software [33] and counting particles in cells using Analyze Particles plugin in Fiji/ImageJ. At least 50 cells per condition were quantified. For the co-localisation between p62 puncta and LC3-GFP-RFP reporter, live cells were imaged on a LSM800 confocal microscope and fluorescence intensity across a line was extracted using Fiji and plotted in Prism 10.1.2 software (GraphPad).

### Mitochondrial clearance

PentaKO HeLa cells transduced with wild-type p62-Halo were transiently transfected with YFP-Parkin, leading to a mixed population of cells expressing YFP-Parkin or not. Mitochondria were cleared from YFP-Parkin expressing cells as in [19] with 10 µM FCCP, substituting CCCP. Briefly, transfection mix contained a 1:1 ratio of YFP-Parkin and blank pcDNA in OptiMEM with Lipofectamine 2000 (11668019, Invitrogen). YFP-Parkin was a gift from Richard Youle (Addgene plasmid # 23955 [34]). Transfection mix was added for 5 hours before being removed and washed with normal media. 24 hours post-transfection, cells were treated with 2 sequential 24-hour treatments of 10 µM FCCP, the media changed again and 48 hours later, cells were treated with Halo ligand (Janelia Fluor 646) ± 40 µM SQ-1 for 5 hours. This was removed and cells treated with 100 nM Mitotracker Green for 30 minutes, with 5 µM Hoechst 33342 added after 15 minutes. Cells were imaged in live-cell using an inverted DMi8 microscope (Leica) with a Plan-Apochromat 63x/1.40 oil immersion lens.

### Assessment of autophagy

For immunoblot analysis, HeLa cells expressing Halo-LC3 and *Npc*^+/+^ and *Npc1^-/-^* MEFs expressing Halo-GFP-LC3 were treated with 50 nM rapamycin, 400 nM bafilomycin A1, 40 µM (HeLa) or 15 µM (MEFs) SQ-1, and 20 µM 7-Bromo-1-heptanol (HaloTag-blocking agent) (H54762, Thermo Scientific Chemicals) [35] for 5 h (HeLa) or 24 h (MEFs), followed by immunoblot analysis using Halo antibody. Autophagic activity was quantified as a ratio of Halo monomer to Halo-LC3 or Halo-GFP-LC3.

For autophagy flux assays using mRFP-GFP-LC3 reporter, cells expressing the reporter were seeded in 35 mm glass bottom dishes and treated for 24 h with 50 nM rapamycin, 400 nM bafilomycin A1, 10 µM SQ-1. For autophagy flux assays in the presence of antioxidants, cells were treated with 40 µM SQ-1 and 1 mM N-Acetyl-L-cysteine, 500 nM MitoQ10, 10 µM S1QEL, 10 µM S3QEL or 10 µM VAS 2870 for 5 h before imaging. Cells were imaged on a LSM700 confocal microscope (Zeiss) and the number of autophagosomes (GFP^+^ RFP^+^ puncta) and autolysosomes (GFP^-^ RFP^+^ puncta) per cell were quantified by outlining single cells as regions of interest using Cellpose software [33] and counting particles in cells using Analyze Particles plugin in Fiji/ImageJ. At least 50 cells per condition were quantified.

### H_2_O_2_ treatments

HeLa cells were seeded in 6-well plates (p62 western blot) or on glass coverslips in 24-well plates (p62 immunofluorescence). After 24 h cells were treated with 30 µM SQ-1 for 18 h followed by H_2_O_2_ at 100 µM or 500 µM for 10 min. Cells were either fixed with 4% PFA for immunofluorescence or protein was extracted for p62 DLCs immunoblots as described below.

### Immunoblot analyses

HeLa and MEFs cells were seeded in 6-well 24 hours prior to treatments. After treatments, cells were washed in ice-cold 1× PBS then lysed in RIPA buffer (150 mM NaCl, 1% NP40, 0.5% NaDoC, 0.1% SDS, 50 mM Tris pH 7.4, supplemented with 1× Halt Protease & Phosphatase inhibitor cocktail (Thermo Scientific) in ddH_2_O) plus 50 mM N-ethylmaleimide. Cell lysates were then centrifuged at 4 °C at 13,000 rpm for 10 min to remove insoluble cellular components. Protein concentration was measured by the DC Protein Assay (BioRad) and a FLUOstar Omega plate reader (BMG Labtech). Samples were prepared by boiling in SDS Loading buffer (BioRad) at 100 °C for 5 min in the presence or absence of 2.5% β-ME (β-mercaptoethanol, Sigma). Amounts equivalent to 20–40 μg of protein were run on 10–12% Tris-Glycine SDS-PAGE gels and transferred to Immobilon-P (Millipore) PVDF membranes using a Trans-Blot SD Semi-Dry Transfer Cell (BioRad). Blots were incubated with a blocking solution (PBS containing 5% fat-free dry milk, 0.1% Tween-20) for 1 h. After washing with PBS, blots were incubated with primary antibodies diluted in blocking solution at 4 °C overnight. Blots were then washed three times for 5 min in PBS with 0.1% Tween-20, incubated with secondary HRP-conjugated α-mouse or α-rabbit (#A2554 and #A0545, Sigma, 1:5000) or α-guinea pig (ab102360, Abcam) antibodies in blocking solution for 45 min at room temperature. Blots were washed three times and incubated for 5 min with the Clarity Western ECL Substrate (BioRad) and signals were detected by chemiluminescence using an iBright Imaging Systems (ThermoFisher Scientific). The following primary antibodies were used: Mouse α-Halo (#G9211, Promega, 1:1000), mouse α-β-actin (STJ96930, St John’s Laboratory, 1:5000), guinea pig α-p62 (#GP62-C, Progen Biotechnik, 1:1000), rabbit α-S6 (2217S, CST, 1:1000), rabbit α-Phospho-p70 S6 Kinase (Thr389) (9205S, CST, 1:1000).

### Mitophagy assay

To enforce mitochondrial respiration, HeLa and MEFs expressing YFP-Parkin and mt-mKeima were cultured for 24 h in galactose medium (glucose-free DMEM (Gibco) supplemented with 10 mM D-galactose, 10 mM HEPES, 1 mM sodium pyruvate, 4 mM L-glutamine, 100 U/mL penicillin/streptomycin and 10% FBS (all from Sigma-Aldrich). After 24 hours, galactose medium was supplemented with 100 nM rapamycin or 10 µM SQ-1 for 24 h. Live-cell mt-mKeima signal was obtained using a DMi8 microscope (Leica) and mitophagy events were determined via following steps using Fiji/ImageJ (version 1.54i; NIH) [18]. Images were masked by applying MaxEntropy threshold algorithm to the images obtained with 561 nm excitation to remove low red signal and background. Within the masks, signals of mt-mKeima were adjusted by applying Enhanced Contrast plugin with saturated = 0.1, normalise, equalise options. Then, images were generated by subtracting the signal at a 480 nm excitation (reporting neutral pH-environment) from the signal at a 561 nm excitation (reporting an acidic pH-environment). Resulting images were binarised with the MaxEntropy threshold algorithm to extract mitolysosomes. The number of puncta per cell was quantified by outlining single cells as regions of interest using Cellpose software [33] and counted using Analyse Particles plugin in Fiji/ImageJ.

### ER-phagy assay

HeLa cells expressing Halo-GFP-KEDL construct were seeded on 35 mm glass bottom dishes. The next day, Halo TMR ligand was added at 1 µM final concentration into the medium supplemented with 100 nM rapamycin or 40 µM SQ-1 for 5 h. The cells were imaged using a DMi8 microscope and ER-phagy events were determined following steps using Fiji/ImageJ. Images were generated by subtracting the signal at a 480 nm excitation (reporting neutral pH-environment) from the signal at a 561 nm excitation (reporting an acidic pH-environment). Resulting images were binarized with the MaxEntropy threshold algorithm to extract ER-phagy events. The number of puncta per cell in images was quantified by outlining single cells as regions of interest using Cellpose software [33] and counted using Analyse Particles plugin in Fiji/ImageJ.

### Immunofluorescence

Fixed cells were permeabilised in methanol for 4 min at -20°C and then blocked for 1 h in 5% normal goat serum (Sigma-Aldrich) in PBS at room temperature. Blocked coverslips were incubated with mouse α-p62 (#610833, BD Sciences, 1:200) overnight at 4°C. Cells were washed three times and incubated with goat α-mouse Alexa Fluor 594 secondary antibody (A-11005, Thermo Fisher Scientific, 1:1000) for 1 h at room temperature. Cells were washed, and coverslips were mounted on slides with Prolong gold antifade with DAPI mounting medium (#P36931, Invitrogen). Fluorescence images were obtained on a Dmi8 microscope (Leica). The number of puncta per cell in images was quantified by outlining single cells as regions of interest using Cellpose software [33] and counted using Analyse Particles plugin in Fiji/ImageJ.

iPSC-derived cortical neurons were washed in PBS, fixed with 4% formaldehyde at room temperature for 15 min, permeabilised with 0.5% Triton X-100 for 10 min (except LC3B staining) or with pre-chilled methanol for 5 min (for LC3B antibody), and incubated with Blocking Buffer (5% goat or donkey serum (Sigma-Aldrich) in PBS with or without (LC3B staining) 0.05% Tween 20 for 1 h at room temperature. Cells were then incubated overnight with primary antibodies at 4 °C, followed by incubation with Alexa Fluor conjugated secondary antibodies for rabbit (A21206, Thermo Fisher Scientific) or mouse (A21203, Thermo Fisher Scientific) for 1 h at room temperature. The coverslips were mounted on glass slides with ProLong Gold antifade reagent with DAPI (Invitrogen). The following primary antibodies were used: rabbit α-LC3B (NB100-2220, Novus Biologicals, 1:200), mouse α-p62 (610832, BD Biosciences, 1:200), mouse α-TUJ1 (4466, CST, 1:200).

### Mitochondrial ROS measurement

*Npc*^+/+^ and *Npc1^-/-^*MEFs seeded in a 35 mm glass bottom dishes were transferred to galactose medium 24 h post seeding. The next day, galactose medium was supplemented with 50 nM rapamycin or 10 µM SQ-1 for 24 h, then stained with 2.5 μM MitoSOX (M36008, Invitrogen) for 10 min, washed three times with galactose medium and imaged on a Dmi8 microscope (Leica). Fluorescence intensity was analysed by outlining single cells as regions of interest and calculation of the integrated density value per cell. The resulting values were then normalised using the MEFs *wt* MitoSOX intensity as a baseline.

### Cytotoxicity assays

Cytotoxicity in *wt*, *Npc1^-/-^*, *Atg5^-/-^* MEFs was measured using Cytotox-Glo Cytotoxicity Assay (G9291, Promega) according to manufacturer instruction. Cells were seeded in a 96-well white plate (Greiner) and switched to galactose medium after 24 h. After 144 h in galactose medium, cells were incubated with Cytotox-Glo Assay Reagent for 15 min at room temperature in the dark, then luminescence was measured using a GloMax plate-reader (Promega) and the readings obtained were attributed to the basal cytotoxicity per well (first reading). To estimate cell population per well, cells were further incubated with Lysis Reagent for 30 min at room temperature in the dark, after which luminescence was measured again (second reading). Population of cell alive was then calculated by subtracting the first reading (basal cytotoxicity per well) to the second reading (indicative of cell population per well) and normalised to the *wt* control condition (DMSO).

TUNEL staining for apoptotic neuronal cells were performed using Click-iT Plus TUNEL Assay kit (C10617, Invitrogen), according to the manufacturer’s protocol. For detection of TUNEL^+^ apoptotic nuclei specifically in neurons, cells were subjected to immunofluorescence by blocking with 3% BSA (in PBS) followed by incubation with anti-TUJ1 antibody (801201, BioLegend) overnight at 4 °C, and thereafter incubated with Alexa Fluor 594 secondary antibody for 1 h at room temperature. Coverslips were mounted on glass slides with ProLong Gold antifade reagent with DAPI (Invitrogen). The quantification of TUNEL^+^ apoptotic nuclei in TUJ1^+^ neuronal cells was performed via fluorescence microscopy, as previously described [23]. The percentage of TUNEL^+^ nuclei was calculated from the total number of TUJ1^+^ cells analysed.

### Modelling of SQ-1-p62 binding

The SQ-1 ligand was docked to the binding site of the p62/SQSTM1 ZZ domain (PDB ID: 5YP8, resolution 1.45 Å) [36]. Piecewise Linear Potential (ChemPLP) [37] and Astex Statistical Potential (ASP) [38] were used in the GOLD (v2024.1) docking algorithm. The docking centre for the binding pocket was defined as the side chain carbonyl carbon of Asp129 (x = 8.242, y = -14.427, z = 197.93) with a radius of 10 Å. Fifty docking runs were allowed for each ligand with default search efficiency (100%). The basic amino acids lysine and arginine were defined as protonated. Furthermore, aspartic and glutamic acids were assumed to be deprotonated.

### DFT calculations

The Gaussian 16 software suite was used with unrestricted DFT. The B3LYP functional hybrid approach was employed [39, 40] and standard 6-31G(d,p) basis set [41, 42] was used for geometry optimisation and frequency analysis (keywords: opt freq). The zero-point vibrational energies (ZPE) were scaled according to Wong (0.9804) [43]. In all cases, normal modes revealed no imaginary frequencies indicating that they represent minima on the potential energy surface. The subsequent energy calculations were then performed with the larger 6-311G(2df, p) basis set. Adiabatic ionisation potentials (IP) and adiabatic electron affinities (EA) were calculated as described [44]. The calculated energies and ZPEs are given in Table S1.

### Statistical analysis

All experiments were carried out in three or more biological replicates from independent cell culture. Graphical data denote the mean ± SEM (of n = 3 biological replicates) and are depicted by column graph scatter dot plot, using Prism 10.1.2 software (GraphPad). Unless indicated otherwise, the *P* values was determined by one-way or two-way ANOVA followed by multiple comparisons using Dunnett’s or Šidák’s statistical hypothesis testing using Prism 10.1.2-10.4.2 software (GraphPad).

## Data Availability

Original images of immunoblotting are provided as supplementary material. All other datasets generated in this study are presented and analysed within this manuscript and are available from the corresponding author upon request.

## Acknowledgements

We are grateful to G. Nelson and Newcastle Bioimaging Unit for imaging support and H. Salmonowicz for graphical design.

## Author contributions

Conceptualisation: J.P., C.S., T.K., W.Y., G.R., J.R., S.S., V.I.K.; Formal analysis: J.P., T.K.; Funding acquisition: J.R., T.K., G.R., V.I.K.; Investigation: J.P., C.S., T.K., E.F., J.R.; Methodology: J.P., C.S., T.K., E.F., J.R., S.S., V.I.K.; Project administration: V.I.K.; Resources: V.I.K.; Supervision: V.I.K.; Visualisation: J.P., C.S., T.K., E.F., J.R.; Writing – original draft: V.I.K.; Writing – review & editing: J.P., T.K., E.F., J.R., W.Y., G.R., S.S, V.I.K.

---

**Supplementary Figure 1.**
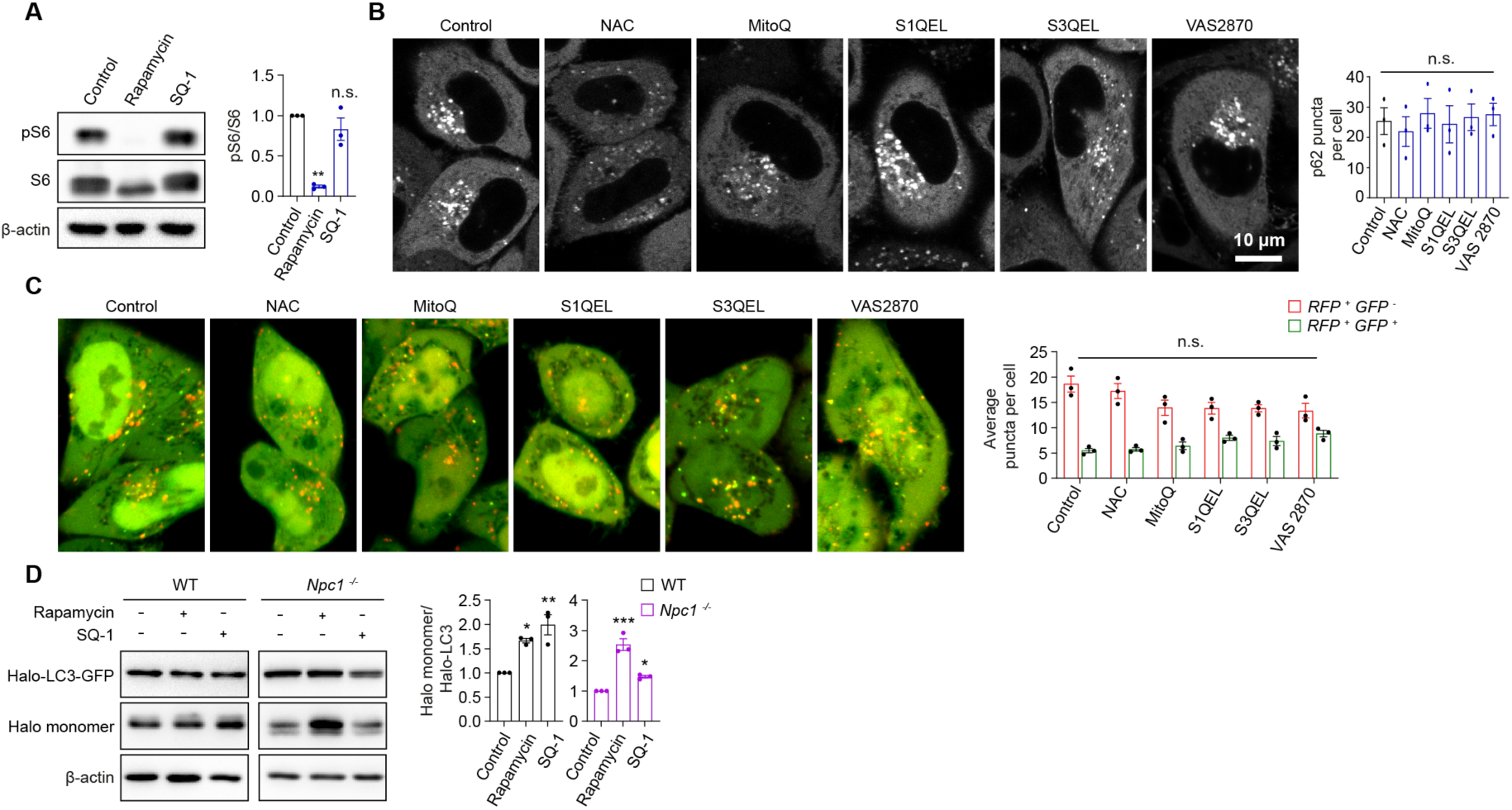
SQ-1 promotes autophagy in an mTORC1-independent manner. **A**, Immunoblot analyses and quantification of S6 and phospho-S6 in HeLa cells treated with 50 nM rapamycin or 10 µM SQ-1 for 24 h. **B**, Confocal fluorescence microscopy images and quantification of p62 puncta in PentaKO HeLa cells expressing Halo-p62 labelled with Halo ligand (Janelia Fluor 646) and treated with 1 mM N-Acetyl-L-cysteine, 500 nM MitoQ10, 10 µM S1QEL, 10 µM S3QEL or 10 µM VAS 2870 for 5 h. **C**, Confocal fluorescence microscopy images and quantification of autophagosomes and autolysosomes in HeLa cells expressing mRFP-GFP tandem fluorescent-tagged LC3 cultured in DMEM supplemented with 1 mM N-Acetyl-L-cysteine, 500 nM MitoQ10, 10 µM S1QEL, 10 µM S3QEL or 10 µM VAS 2870 for 5 h. **D**, Immunoblot analyses and quantification of Halo monomer accumulation in *Npc1*^+/+^ and *Npc1*^-/-^ MEFs expressing Halo-LC3 cultured for 24 h in galactose medium supplemented with 50 nM rapamycin or 15 µM SQ-1. Data are mean ± SEM of n = 3 biological replicates. P values were calculated by one-way ANOVA followed by multiple comparisons with Dunnett’s test (A, B) or two-way ANOVA followed by multiple comparisons with Šidák’s test (C). *P< 0.05; **P<0.01; ***P<0.001; ns (non-significant).

**Table S1.** The single point and corrected zero-point vibrational energies (ZPE) of SQ-1 in hartrees (a.u.).

| Nr. | Systems | Energy/a.u. | ZPE/a.u. <sup>a</sup> |
| --- | --- | --- | --- |
| Parent | SQ-1 | -2120.714688 | 0.471317496 |
|  | IP | -2120.436381 | 0.469237087 |
|  | EA | -2120.751507 | 0.465686078 |

<sup>a</sup> Corrected according to [43].

